# Dynamic analysis and control of a rice-pest system under transcritical bifurcations

**DOI:** 10.1101/2021.09.07.459216

**Authors:** Sajib Mandal, Sebastian Oberst, Md. Haider Ali Biswas, Md. Sirajul Islam

## Abstract

Cultural methods were widely applied at an early stage of agricultural pest management but then replaced over time through pesticides with adverse effects on crop quality and the environment due to extensive and long-term use. In this study, we have reformulated a rice-pest-control model by first modelling a rice-pest system rigorously to then extend it to become an optimal control problem. We consider both, cultural methods and pesticides, and minimize thereby the density of agricultural pests, to increase the production of rice and to reduce gross annual losses. Pesticides have been applied only in an emergency scenario, which reduces environmental pollution and damage to nearby ecosystems. For the emergency case, we have developed a decision model to mitigate potential risks. The formulated models are verified by stability analysis at equilibrium points and investigated through transcritical bifurcations. Moreover, we have extensively confirmed our main results by numerical investigations and discussed the biological implications in more detail.

## 1. Introduction

The production of rice increases annually along with a growing human world population. Improper irrigation, environmental or climatic factors, and the overuse of pesticides [1], hamper its production significantly [2]. Due to high yield losses, costly research and development, and their application of disease and pest resistant crops has become increasingly popular in recent years [3]. Agricultural pests are a driving factor behind those efforts. Animals (insects, birds, rodents), pathogens, or weeds are commonly considered as negative factors. Without pest control, the combined annual losses of rice would exceed 70% [4] mostly due to infestations of flies of the species *Gonatocerus* spp. (Mymaridae), or *Megaselia* spp. (Phoridae) [5]. Yet, despite applying about three billion tons of pesticides annually worldwide, up to 40% of the total rice yield is lost [6,7].

### Control methods

To avoid large and unforeseen losses, agricultural pests need to be controlled. Choosing the best variety of local growing conditions and maintaining a healthy crop is the first line of defence. The first prevention strategy then usually consists of applying cultural methods [8]. Cultural methods e.g., crop rotation, soil enhancement, removal of diseased plants and proper irrigation do not require chemical substances, are less invasive and naturally contribute to increased productivity [9]. Pest species of certain crops decline due to the deficiency of food when a different crop species becomes the focus of cultivation. On the other hand, when plants have enough space so that the sunlight can reach the ground some insect pests can be decreased by more than 96% [10] facilitating visibility to predatory species [11].

Conventionally, chemical controls are used in modern agriculture. Many chemicals such as neonicotinoids or glyphosate are applied annually [2] which, however, are increasingly under scrutiny due to their long-lasting negative effects on the environment, especially the insect populations – essential for pollinating more than 80% of the crop [12]. Other chemical controls include the use of DDT, BHC, 2-D, or 4-D, etc. [2] which have been already banned in many countries. Because of the problems with chemicals alternative controls have been developed including e.g. cultural methods such as field sanitation, crop rotation, tillage, etc., biological controls, or combinations thereof [3,9]. Biological controls constitute effective methods including natural enemies of insect pests as control agents such as parasitoids (e.g., Diapriidae for flies) [13], predators (e.g., *Ophionea nigrofasciata*) [14], genetic sterilisation [15], pathogens (viral infections) [16], or the exploitation of competitor relationships (e.g., a competition for preying between the protist *Didinium* and *Paramecium*) [17]. Further, the application of artificial signals such as that mimicking a female pheromone can be applied to confuse heterospecifics or to trap reproductives [18] an alternative with no harmful chemicals being emitted [14].

Optimised integrated pest management (IPM) strategies often consist of a set of impulsive differential equations (IDEs) [19]. Sun et al. [3] formulated an optimal control strategy approaching a predator-prey model considering biological and chemical control variables and indicated that the chemical control acts faster. By incorporating different regulatory methods, the *model interplay*, itself between pesticides and natural enemies including factors such as residual effects and spray times of pesticides, or release time of natural enemies has shown to have a significant impact on pest species populations [20]. Biological controls imply using mathematical predator-prey models with a periodic release of predators [21], periodic application of pesticides to the pest species [22], and pests infected by bacterial diseases or viruses [23]. The Lotka-Volterra system as a common mathematical predator-prey model consists of a pair of first-order nonlinear ordinary differential equations (NODEs) [24]. However, biological controls are expensive to impose and may require a long-term process and a pest density at a maintainable threshold [25]. Therefore, IPM strategies focusing only on biological controls are limited and usually require several input factors such as host-parasite ratios, starting and ending point population densities, parasite population or insecticide dosage as well as release duration, and levels of parasitism and host-feeding. Whilst previous works mainly focused on predator-prey relationships and pest control, there is limited research on rice pest management. In those works, there are some interesting questions such as which control technique between natural methods and pesticides is more effective in rice pests management and which can also increase annual production of rice has not been described properly. In particular, how and when pesticides should be applied so that adverse effects of pesticides on the environment can be avoided, have not been addressed in more detail.

Thus, in the present study, a rice-pest-control model describing *effective control techniques* and a decision model describing *proper application strategies with minimum adversity of pesticides on the environment* are investigated. Firstly, we formulate a novel rice-pest-control model based on an already developed rice-pest model [Eq. (S7), presented in Supplementary Material A.3] by setting up the interrelationship between the production of rice and the corresponding pests. Here, we adopt two control techniques, cultural methods and pesticides. We then extend the rice-pest-control model to become an optimal control problem aiming to reduce the annual losses and to increase the annual production of rice by incorporating chemicals with the purpose to evaluate a hybrid cultural/chemical IPM strategy. Secondly, when the density of pests in the cultivation becomes so high that it could not possibly be anymore controlled by cultural methods, pesticides as a last resort of controlling the agricultural pests are applied. We develop a decision model for the first time, which decides when the pesticides should be applied or stopped, to control the adverse effects of the long-term and extensive use of pesticides on the environment and nearby ecosystems. The rice-pest-control model is verified by analysis and obtains the necessary conditions for the optimality applying *Pontryagin’s maximum principle* in terms of the *Hamiltonian H*(*t*), whereas the rice-pest model is verified by analysis, experienced stability analysis at equilibrium points, and investigated by transcritical bifurcation analysis.

## 2. Dynamical modelling in form of a hybrid natural/chemical IPM strategy

### 2.1 Assumption and formulation of the optimal control problem

Millions of metric tons (MMT) of rice are lost annually only due to pest infestations amounting to about 37% of the annual production in the world. With the increase in the annual rice yield, the number of annual production and losses gradually increases as shown in Fig. 1. Fig. 1 depicts the annual losses from 1960 (~10.6 MMT) to 2013 (~41.05 MMT). On the other hand, the mathematical model of the rice-pest system, expressed by Eq. (S7) that presented in Supplementary Material A.3, shows that the production of rice inversely changes with the pest population density.

**Fig. 1.**
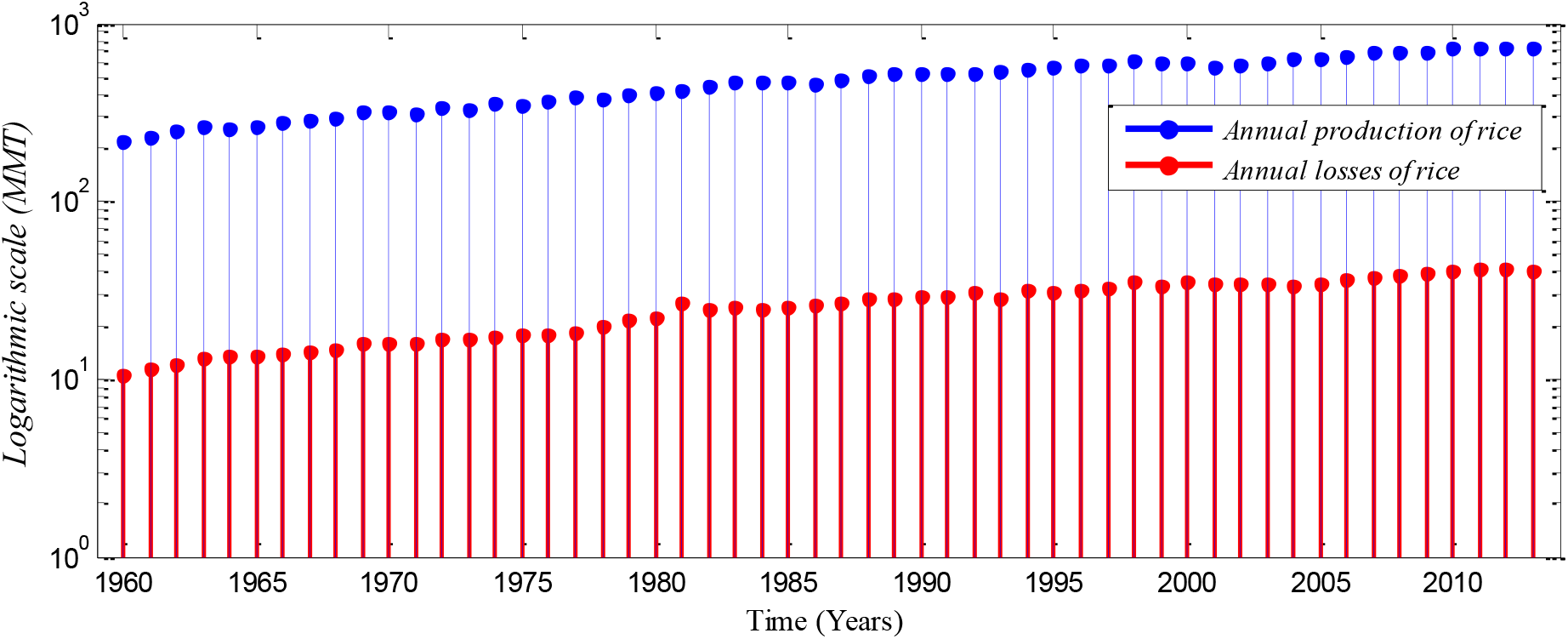
Annual production and losses (due to the attack of pests) of rice from 1960-2013 [26] represented under a logarithmic scale with MMT measurement unit. In comparison to the production, we see that annual losses increased over the years and their absolute number is very high.

**Fig. 2:**
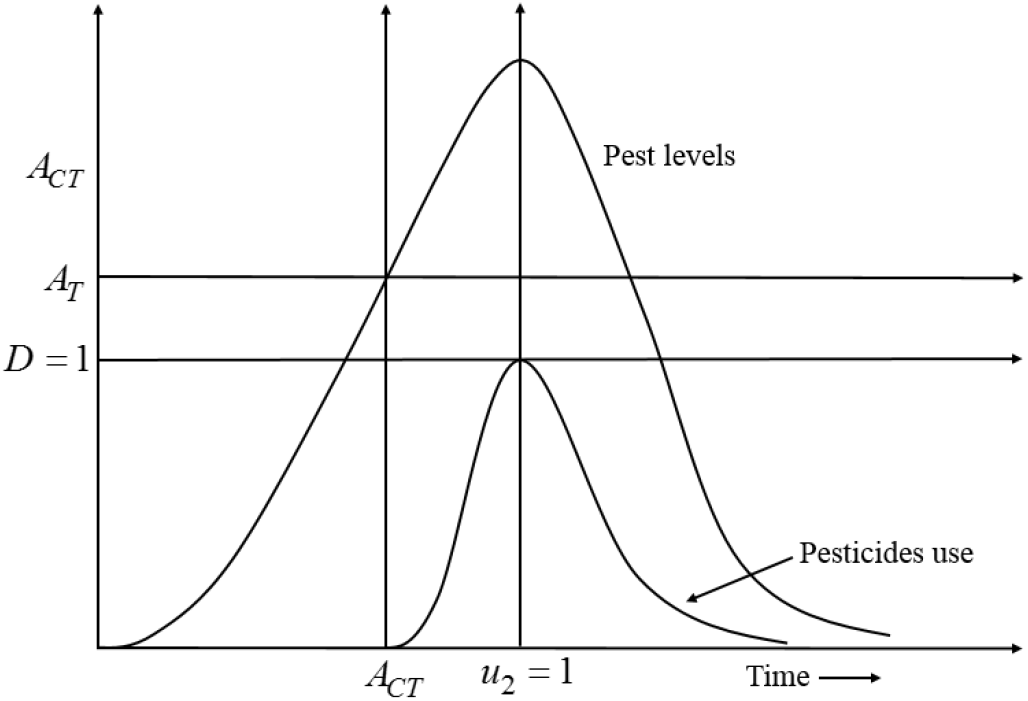
A decision-making diagram of the model expressed by Eq. (1) describing the best time of pesticides application. Here, *D* = 1 represents the acceptable damage threshold reaches the maximum level, which means environmental pollution reaches the most acceptable level; *A_T_* denotes the action threshold to determine the use of pesticides to control the pest population; *A_CT_* represents the interval in which pesticides should be applied.

Cultural methods consist of some methods or techniques that arise crop production naturally by controlling pests. Good soil preparation, use of indigenous varieties, use of mesh screen, pruning, encouraging insect predators, and crop rotation are the most used cultural methods [27]. Chemical control is the most effective strategy for instantaneous pest control when cultural methods cannot be used. However, the overuse of pesticides and insecticides seriously hampers the food quality and adversely impacts aquatic ecosystems [28].

Wiping out the entire pest population from the agricultural field is impossible, and an attempt can be unsafe, expensive, and may lead to a rebound of pest numbers [8]. To control the pest population, we first apply cultural methods that are safe, cheap, and easily applicable. We then model the application of chemical controls (pesticides) only when the unacceptable action thresholds are crossed [8]. The application of pesticides depends on the species of pest and their density.

An important question is the determination of the point in time when to apply pesticides and for how long it should continue. Let *u*_1_ denote cultural methods which are used incessantly throughout the entire cultivation period because they are natural and safe. Let *u*_2_ denote chemical controls (pesticides) which are used only when the pest level crosses the acceptable threshold. The controls *u*_1_ and *u*_2_ both take values between 0 and 1. Here, *u*_1_ *u*_2_ = 0 denote that no control is applied and *u*_1_ *u*_2_ = 1 indicate controls are applied with full effort. Let *A_T_* denote the unacceptable threshold (or action threshold) of the pest population. The unacceptable threshold can be defined as: *A_T_* = *μ*Δ, where *μ* denotes an unacceptable pest population per unit area and Δ represents the total area of a field. The threshold can depend on the pest variety, pest size, and site/region [8]. Let *D*(*x*_2_, *u*_2_, *E*) denote the function of the nearest ecosystem which is potentially subject to pesticide damage, with *x_2_* being the pest population and *E* represents environmental damage. *E* is defined in the interval [0, 1] with *E* = 1 denoting that the pollution reaches the most acceptable level (*E* = 0 denotes zero pollution). The damage function, *D*, is increased in the rise of *x*_2_ and *E* but decreased in the application of *u*_2_, with *D* ∈ [0,1] [29,30]. *D* = 1 represents the upper level of acceptable damage threshold, meaning the application of pesticides should be stopped i.e., *u*_2_ = 0. Conversely, *D* = 0 denotes that there is no damage, meaning pesticides can be applied with full effort i.e., *u*_2_ = 1. Here *E* is proportionately increased with the application of *u*_2_. If *E* → 1 as *u*_2_ → 1 which leads *D* → 1 that means the use of pesticides should be stopped immediately. To reflect this, the following decision model has been developed:

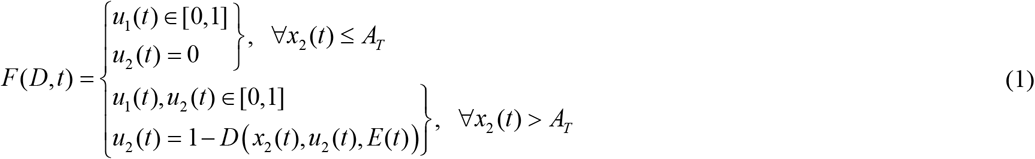

A control model has been proposed on a developed rice-pest model, expressed by Eq. (S7) that presented in Supplementary Material A.3, where cultural methods and chemical controls are applied as two control variables with the aim to minimize the density of rice pests. The dynamical relationships between the annual production of rice and the pest population under control are described below:

i. When cultural methods are applied during cultivation, the production rate of rice increases rapidly. Since the use of cultural methods does not depend on the density of rice pests, let *u*_1_ *x*_1_ be the increment in the annual production of rice. Therefore, the first equation of Eq. (S7), presented in Supplementary Material A.3, can be represented by including control as

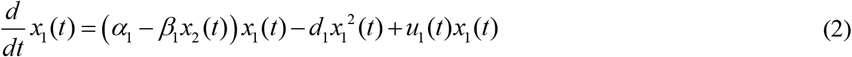

here, *x*_1_(*t*) represents the annual production of rice, *α*_1_ shows the reproduction rate of rice, *β*_1_ represents the loss rate of *x*_1_(*t*) due to the consumption of pests, *d_1_* presents the decrease rate due to intraspecific competition in species *x*_1_(*t*) due to natural causes that are not related to pests e.g., viral infections, droughts or floods.
ii. When cultural methods are applied in rice cultivation, several rice pests die off because of soil rotation and the presence of predators. Therefore, the cultural methods decline the density of rice pests and let *u*_1_*x*_1_*x*_2_ be the declining number of pests due to the adoption of cultural methods. On the other hand, when emergency situations, pesticides are applied according to Eq. (1), which significantly reduces the pest population. Since the application of pesticides in the cultivation of rice depends on the amount of cultivated area, let *u*_2_*x*_1_*x*_2_ be the decline in the density of pest population after the use of pesticides. Hence, the second equation of Eq. (S7) can be represented considering the decision model (1) as in the following

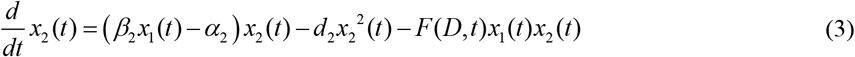

here, *x*_2_(*t*) represents the density of rice pests at time *t, β*_2_ shows the energy gain rate of pest population by consuming rice, *α*_2_ represents the decline rate of the pest’s population proportionally with the decline of rice production, *d*_2_ shows the intraspecific competition between *x*_2_(*t*) due to natural causes that not related to *x*_1_(*t*), e.g. viral infection and heavy rains.

The modified rice-pest system (S7) under all controls can be represented by arranging Eq. (2) and Eq. (3) as in the following:

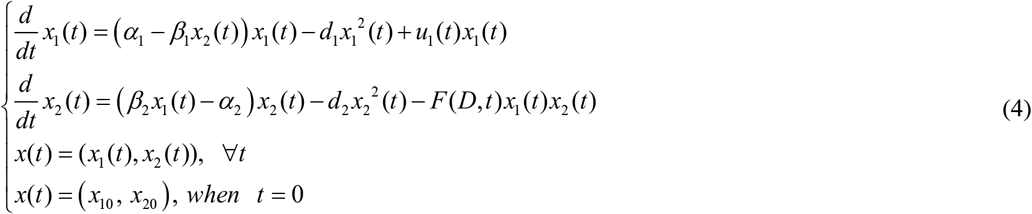

For more details, please look at Supplementary Material A.3.

The characteristics of the controls are represented in the following measurable control set

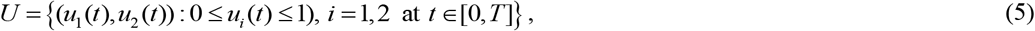

where *T* is a preselected period for applied controls. The objective functional of the control model (4) becomes

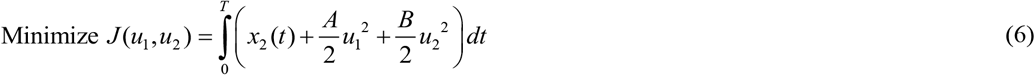

The optimal control model which approximates model (4) can be represented [31] as:

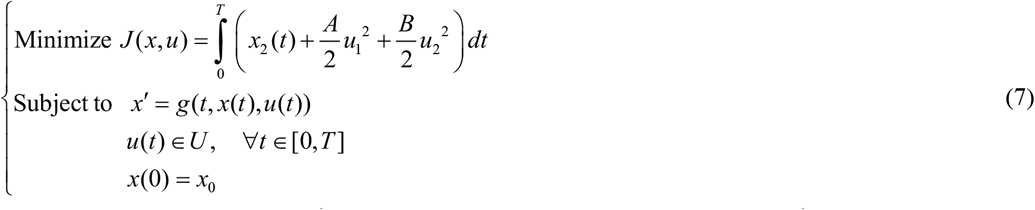

where 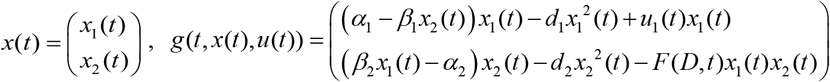 and 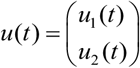 *A* and *B* are used for cost balancing weight parameters for the control variables *u*_1_ and *u*_2_, respectively; the function *g* is continuously differentiable; and functions *u*(*t*) and *x*(*t*) are piecewise continuous differentiable. In this problem, *u*(*t*) belongs to a certain space *U* that may be a piecewise continuous function or a space of measurable functions which satisfy all constraints of the problem. The main goal of the objective functional is to increase the annual rice yield by minimizing the pest population by simultaneously considering the controls with the lowest costs. A schematic diagram of the rice-pest-control system (4) is shown in Fig. 3 to illustrate the dynamic behaviour of the species under control.

**Fig. 3.**
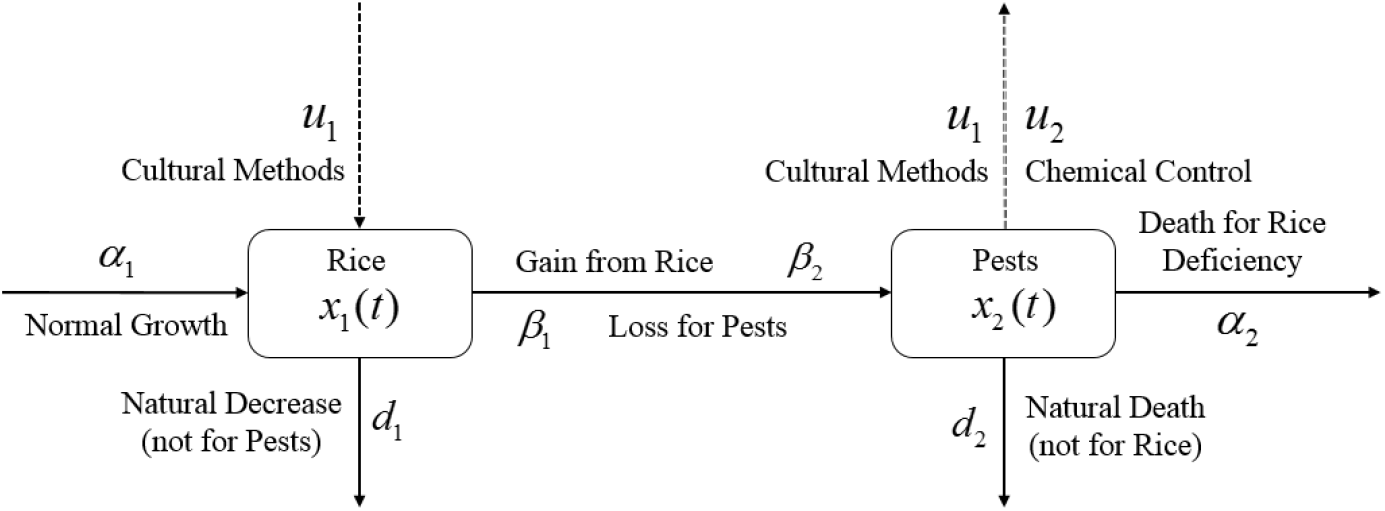
Schematic diagram of the rice-pest-control system (4) describes the rice-pest system (S7) under control. The diagram also shows that the control strategies, cultural methods and chemical control (pesticides), increase the production of rice and decline the corresponding pest population.

### 2.2 Characterization of the optimal control

To estimate the necessary conditions for the optimality of the optimal control problem (7), *Pontryagin’s maximum principle* has been imposed in terms of the *Hamiltonian H*(*t*) defined as [31]

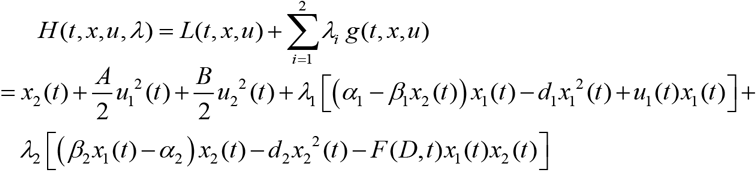

where *λ_i_, i* = 1,2 is the co-state variable which satisfies the following adjoint equations,

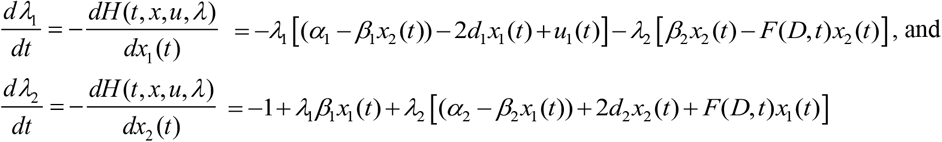

as well as the *transversality conditions λ*_1_(*T*) = 0 and *λ*_2_(*T*) = 0.

Now, to obtain the optimal solution of the controls, Theorem 2.1, and Theorem 2.2 must be proven as shown below by applying *Pontryagin’s maximum principle*.

#### Theorem 2.1.

*The control variables for the acceptable damage threshold attain the optimal solutions* 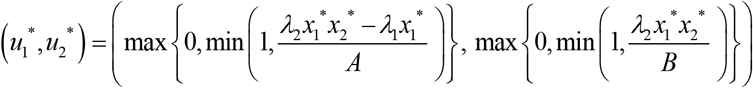 *for which the objective function J over U is minimized*.

**Proof:** At the acceptable damage threshold, there is no use of pesticides i.e., *u*_2_ = 0. Therefore, let’s differentiate the *Hamiltonian* (*H*) with respect to the control variable *u*_1_ only, then it becomes as

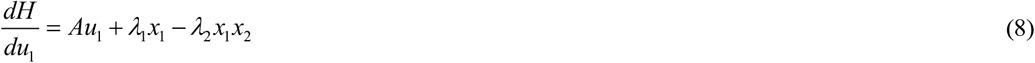

By applying the conditions of optimality in Eq. (8), the characterization of the control variable *u*_1_

i. when 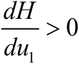 then 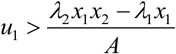 but for the minimization problem *u*_1_ = 0
ii. when 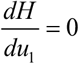 then 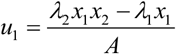
iii. when 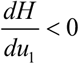 then 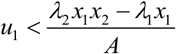 but for the minimization problem *u*_1_ = 1

Therefore 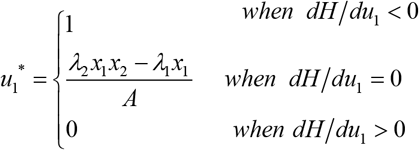, which can be written in the following compact form

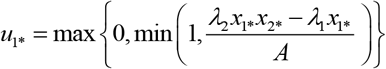

Since *u*_2_ = 0 at the acceptable threshold, the compact form of *u*_2_ is *u*_2*_ = 0. Then, the optimal solutions of the control variables for the acceptable threshold are

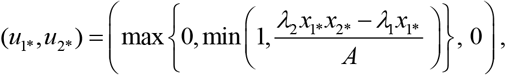

hence, it completes the proof.

#### Theorem 2.2

*The control variables for the action threshold attain the optimal solutions* 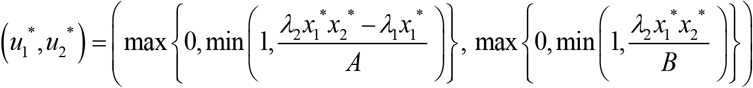 *for which the objective function J over U is minimized*.

**Proof:** For the action threshold, the decision model is fully active i.e., *F*(*D,t*) = *u*_1_(*t*) + *u*_2_(*t*). Therefore, let’s differentiate the *Hamiltonian* (*H*) with respect to the control variables *u*_1_ and *u*_2_, then it becomes as

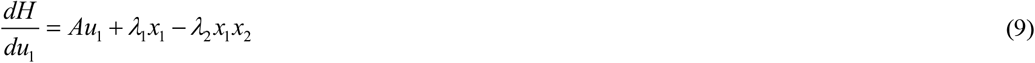

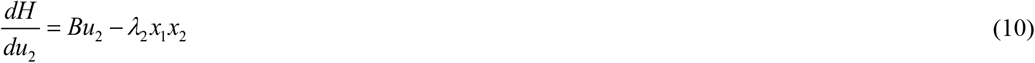

By applying the conditions of optimality in Eq. (9), the characterization of the control variable *u*_1_

i. when 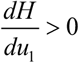 then 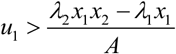 but for the minimization problem *u*_1_ = 0
ii. when 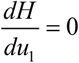 then 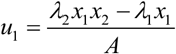
iii. when 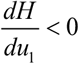 then 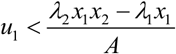 but for the minimization problem *u*_1_ = 1

Therefore 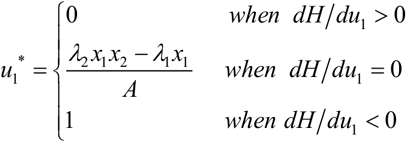, which can be written in the following compact form

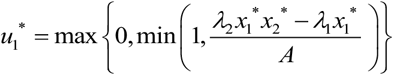

Similarly, by applying the conditions of optimality in Eq. (10), the characterization of the control variable *u*_2_ becomes 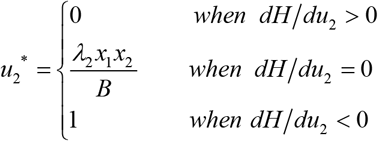, which can be written in the following compact form

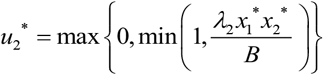

Then, the optimal solutions of the control variables are

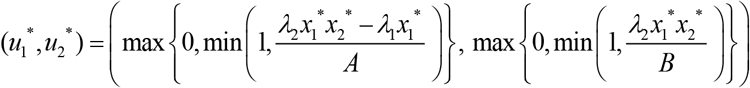

Hence the theorem completes the proof.

## 3. Results

Numerical simulations have been carried out for the rice-pest model (S7) and rice-pest-control model (4) using the ode45 solver in MATLAB. This section aims to illustrate the dynamic change of the annual production of rice and the growth of the rice pests before and after adopting the control methods. In this case, the numerical values of parameters used in the simulations are taken from Table 1 and the initial conditions of the state variables chosen are *x*_10_ = 4.679, *x*_20_ = 0.05085. We consider 12 months (1 year) for the simulations.

**Table 1.**
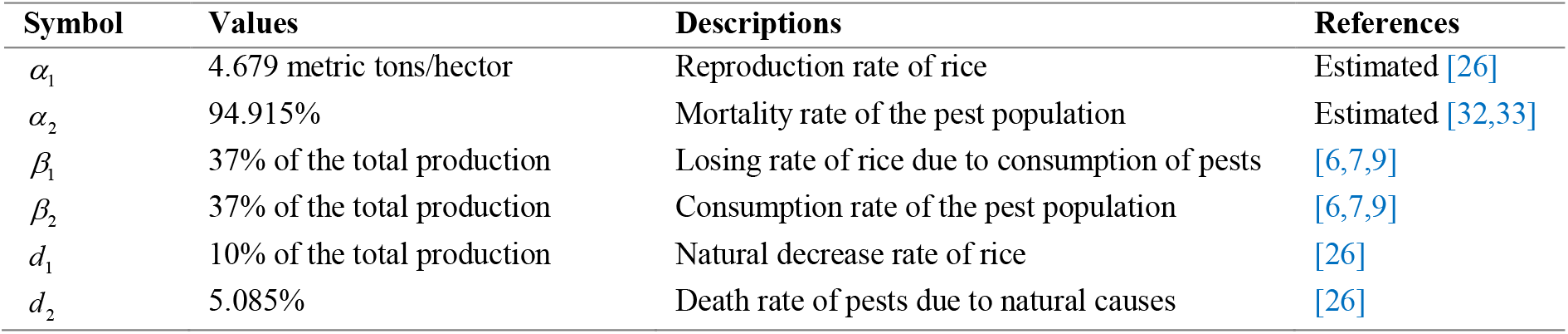
A description of parameters used in this study, including the numerical value. All of these are secondary parameters (derived/estimated or collected from other sources), two of which are estimated. We have conducted statistical analysis for parametric estimation after collecting and observing the corresponding data collected from different research [26,32,33]. For more details, please look at Supplementary Material C.

Both the time series and phase portrait of the considered dynamics are shown in Fig. 4. Initially, the growth of the rice plants sharply increases due to low numbers in the pest population. When the pest population increases by getting sufficient food, the growth of the rice plants is hampered so that eventually the pest population subdues (which in turn allows rice to grow again) [4,8]. The phase portrait is shown in Fig. 4 (b) spiralling out to a stable equilibrium at 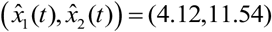.

**Fig. 4.**
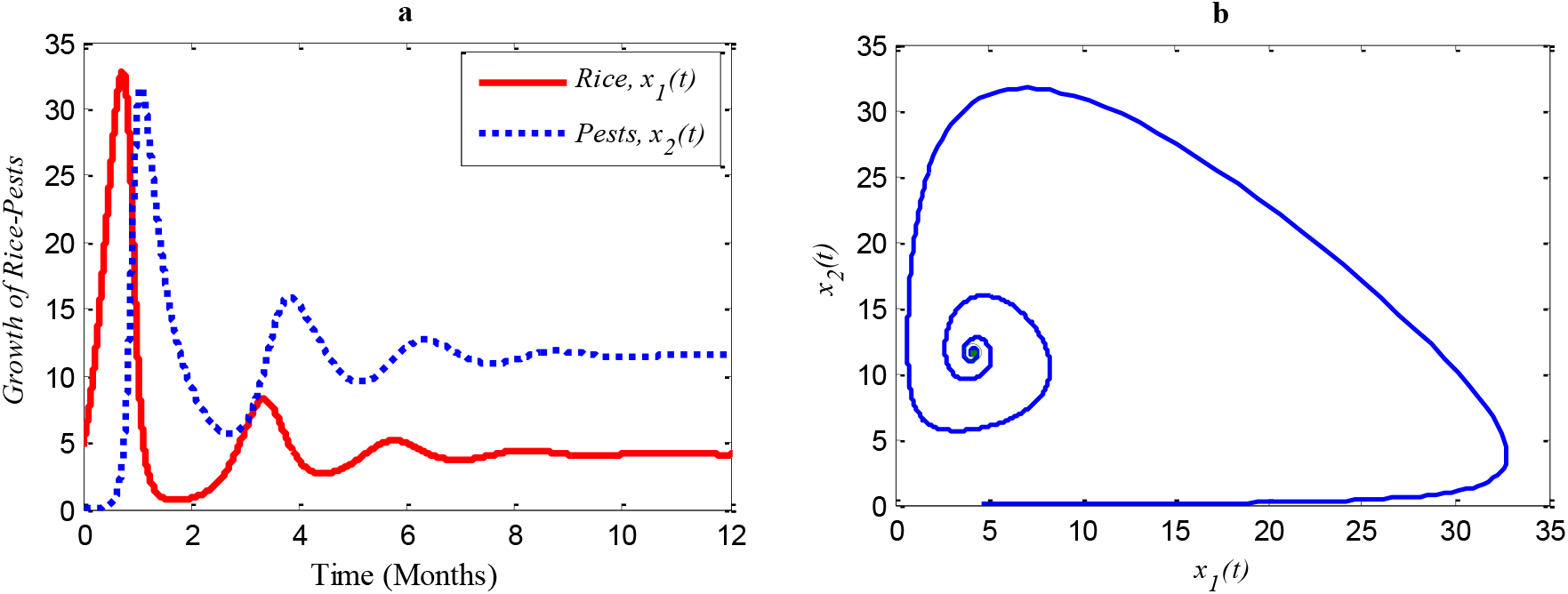
Time series of (a) Rice-pest population, and (b) phase portrait of the rice-pest system (S7) with the solution 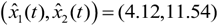

Fig. 5 (a) and (b) show (i) when the consumption rate *β*_2_ of the pest population is reduced from 37% to 25%, the density of pest population decreases so that the rice is increasing again [4,8]. Similarly, in (ii): when the consumption rate declines from 25% to 15%, the density of pests declines sharper than before, and the rice grows better again [4,8].

**Fig. 5.**
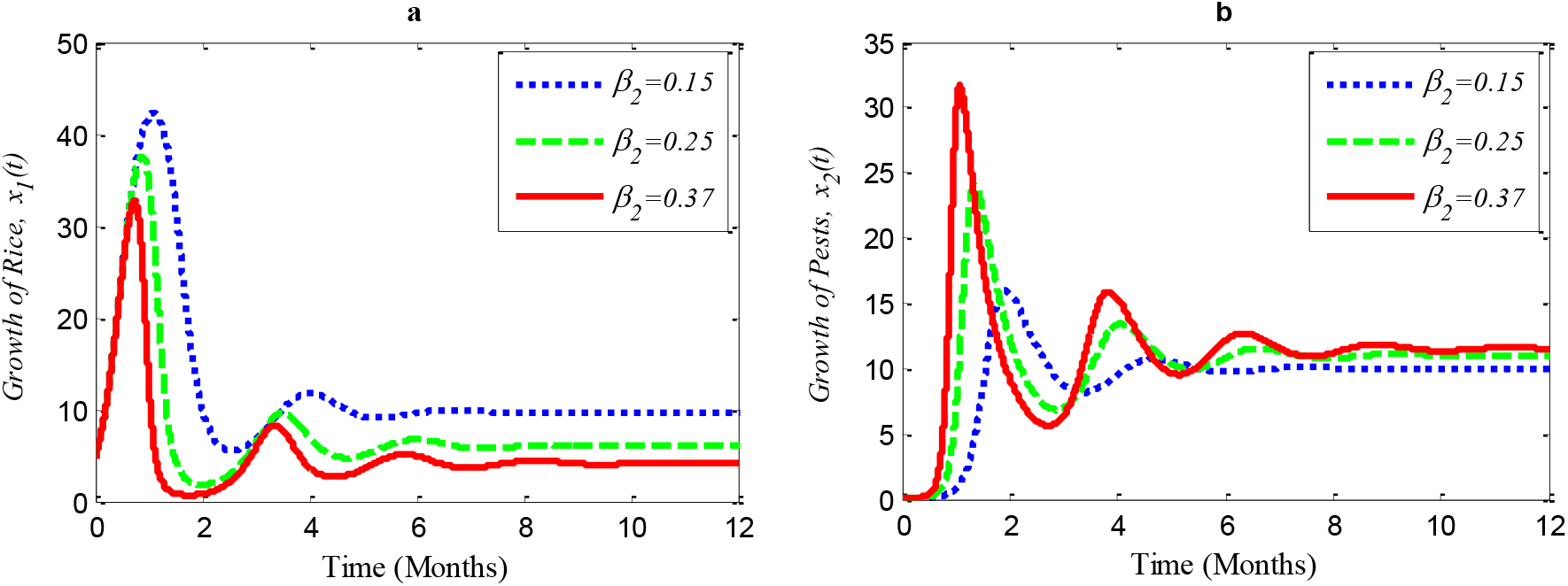
Time series of (a) annual rice production, and (b) rice pest population for different consumption rates of the pests.

According to the result of Fig. 4 (a), the annual production of rice approaches 4.12 Metric tons/hector (Mt/h) whereas the growth rate of pests approaches 11.54%. The changes in the growth of rice for different consumption rates of pests are described in Fig. 5 which let us conclude that the production of rice can be increased if the consumption rate of pests is controlled.

When controls are applied, the results change. The results of the rice-pest-control system (4) are described for the following three scenarios designed on the efficacy of the control variable *u_1_* (cultural methods) and *u*_2_ (chemical controls): (i) *u*_1_ ≠ 0 and *u*_2_ = 0, (ii) *u*_1_ = 0 and *u*_2_ ≠ 0, (iii) *u*_1_ ≠ 0 and *u*_2_ ≠ 0. The simulations are performed distinguishing the cases “without control” and “with control”. Here, “without control” represents the results of the rice-pest system (S7) meaning there is no control strategy (*u*_1_ = 0, *u*_2_ = 0).

i. When only the cultural methods are implemented to the system as a control variable, the annual losses of rice decline from 41.05 to 38.39 MMT approximately, at the same time, the production rate of rice increases from 4.12 to 8.81 Mt/h approximately [8–10] which are shown in Fig. 6 (a). As a result of adopting cultural methods, the density of rice pests declines from 11.54% to 10.79% approximately over one period due to e.g., the enhancement of soil and rotation of crops [8–10] as shown in Fig. 6 (b). Fig. 6 (c) shows that the system under the application of cultural methods converges to the stable equilibrium point at 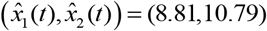.
ii. When only the chemical control is implemented according to the condition expressed in Eq. (1), the pests population comes down from 11.54% to 9.17% approximately that’s just over one-fifth of the total population [2,12] shown in Fig. 7 (b). Because of the decreasing pest population, the annual losses of rice drop sharply from 41.05 to 32.62 MMT approximately with a decrease approximating 25%. As a result, the annual rice production rate substantially grows from 4.12 to 12.86 Mt/h approximately [2,12] which is shown in Fig. 7 (a). The system converges to a stable equilibrium point at 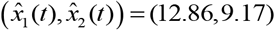, presented in Fig. 7 (c).
iii. When the cultural methods are used continuously, and chemical controls are applied only in the emergency (Eq. (1)) the density of the pest population decreases significantly from 33.02% to 7.73% (Fig. 8 (b)) [2,8–10]. As a result, the annual losses of rice dramatically fall from about 41.05 to 27.49 MMT. After simultaneously applying controls the annual rice production increases considerably and reaches 19.18 Mt/h [2,8–10] as presented in Fig. 8 (a) thereby approaching the equilibrium point 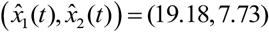 as shown in Fig. 8 (c).

**Fig. 6.**
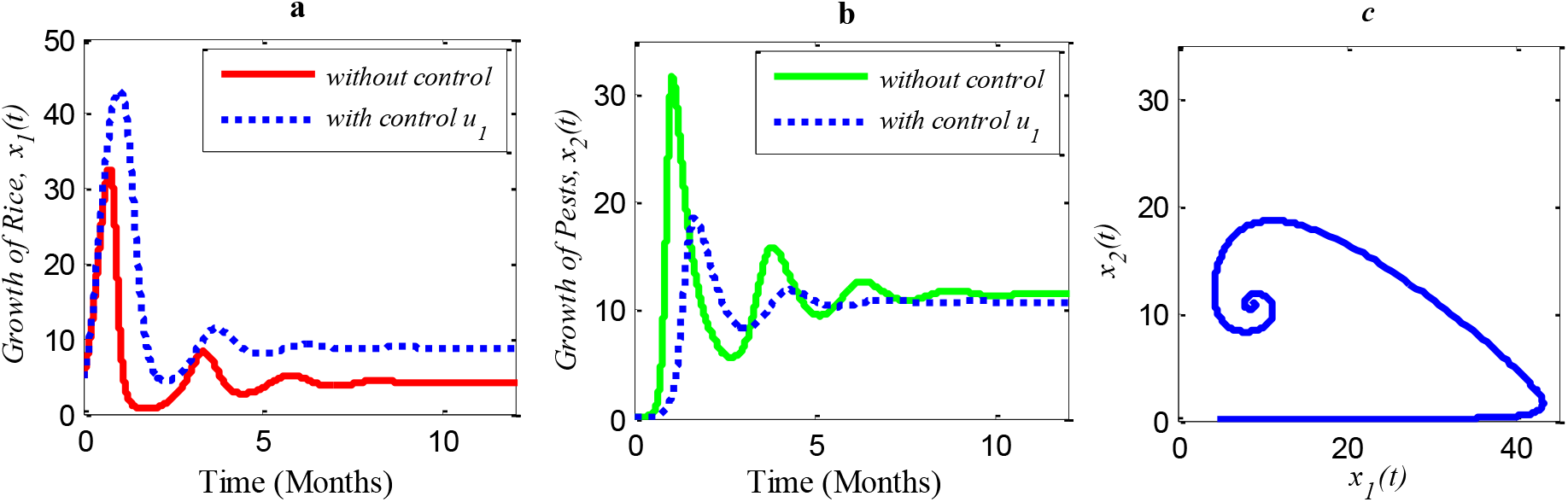
Time series of (a) annual rice production under control *u*_1_ only, (b) pests population under control *u*_1_ only, (c) phase portrait of the rice-pest-control system (4) when only *u*_1_ is adopted, where the solution is 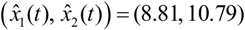.

**Fig. 7.**
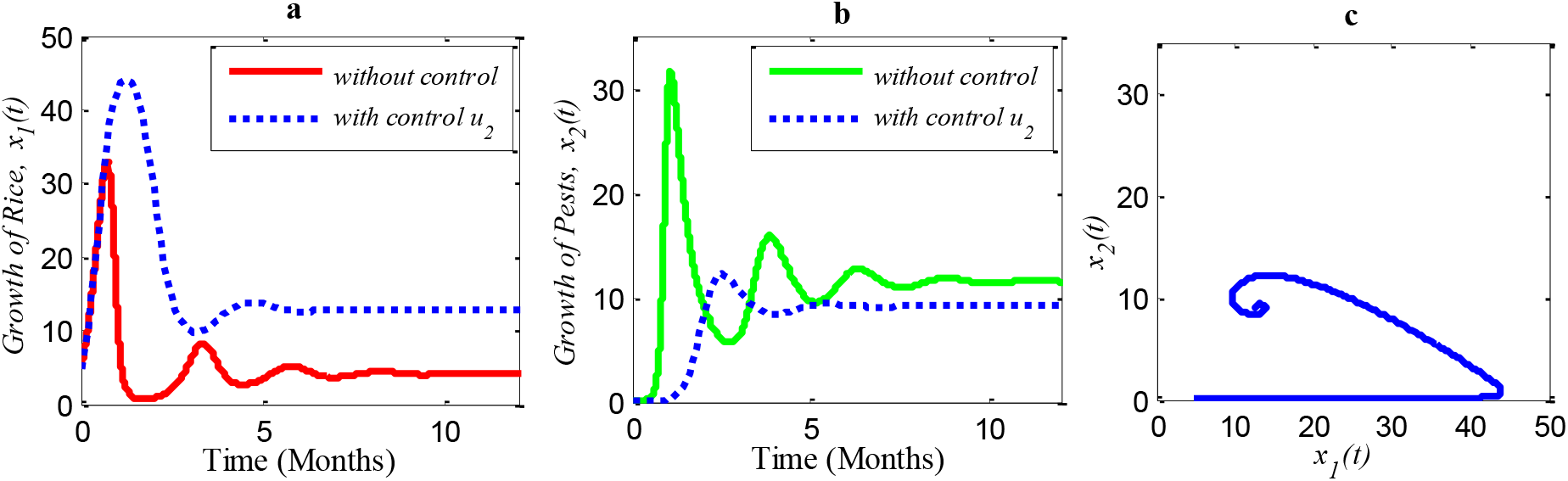
Time series of (a) annual production of rice under control *u*_2_ only, (b) pests population under control *u*_2_ only, (c) phase portrait of the rice-pest-control system when only *u*_2_ is adopted and the solution is 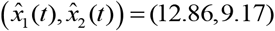.

**Fig. 8.**
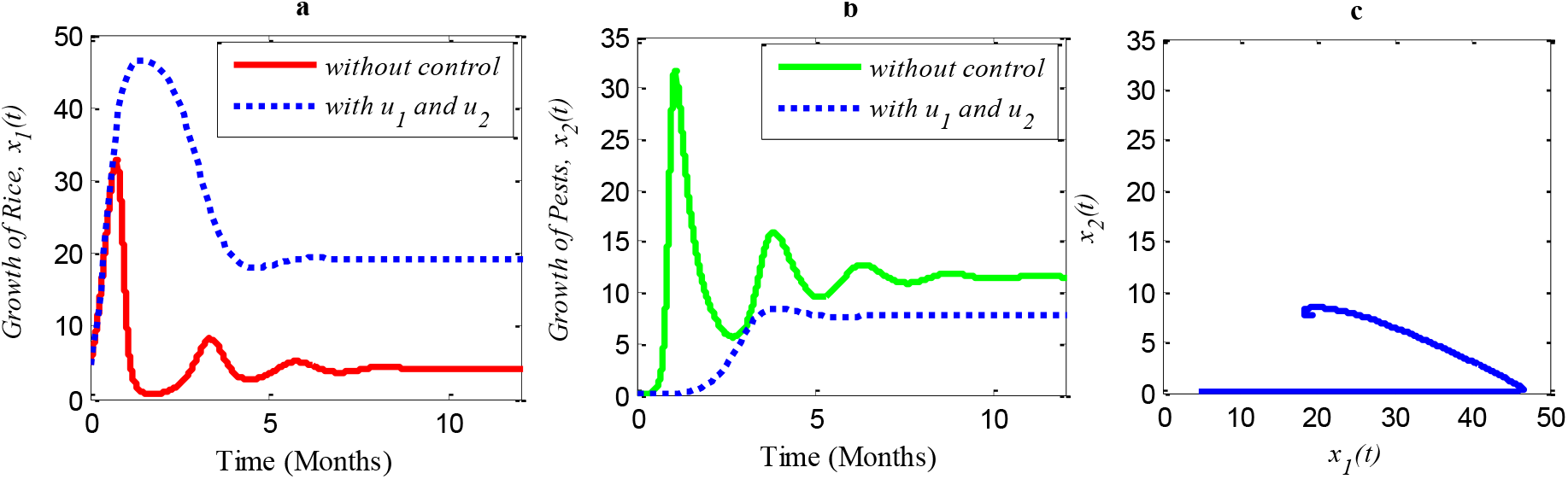
Time series of (a) annual production of rice under controls *u*_1_ and *u*_2_, (b) pests population under controls *u*_1_ and *u_2_*, (c) phase portrait of the rice-pest-control system when both controls *u*_1_ and *u_2_* are implemented, where the solution is 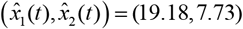

Next, we made a numerical comparison to analyse the results of all situations. The dynamic changes in the growth of the state variables in the three scenarios are shown in Fig. 9. As can be seen from Fig. 9, the annual production of rice increases significantly under both control strategies instead of just one, in contrast, the rice pests decrease dramatically. It also shows that cultural methods can control the density of the pest population, but chemical controls are comparatively more effective. From scenarios (i) to (iii), it is concluded that scenario (iii) is the best strategy to increase the annual rice production and reduce the density of rice pests.

**Fig. 9.**
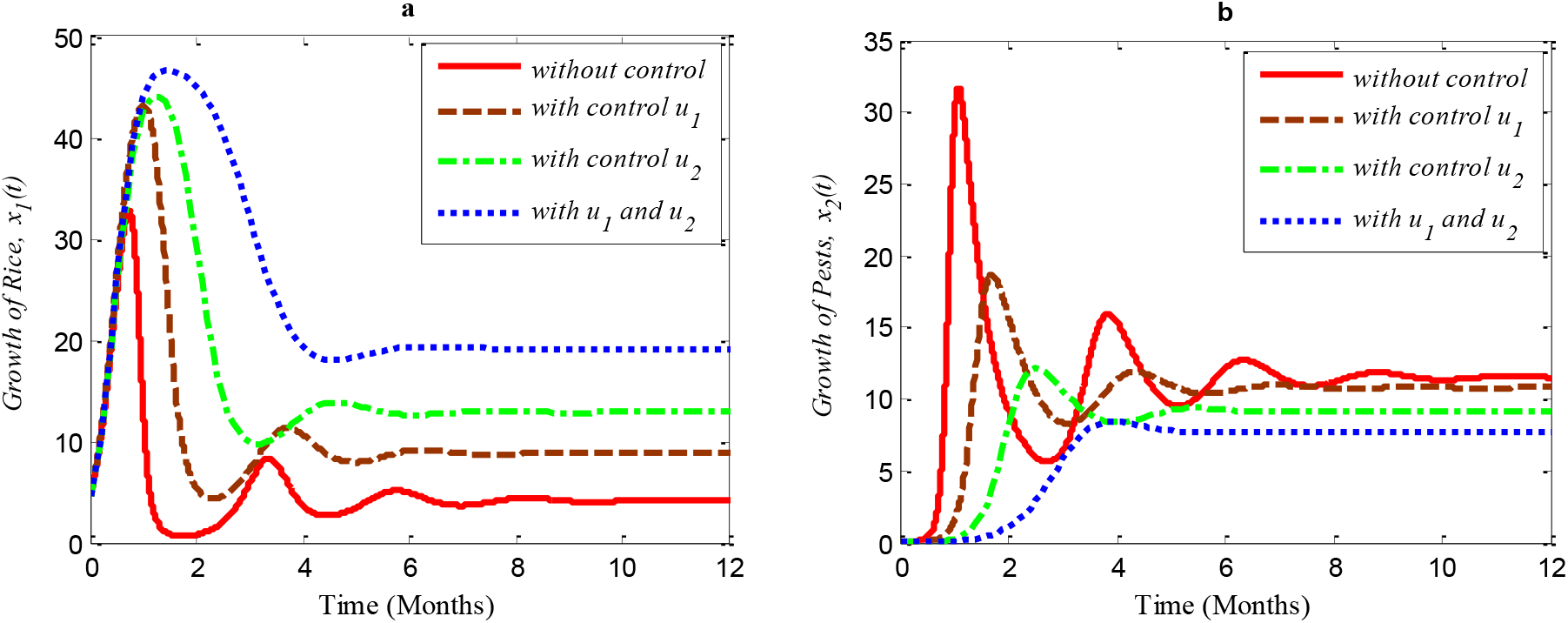
A comparison between the scenarios (i) to (iii). Here, (a) represents the time series of the annual production of rice, and (b) shows the time series of pest populations under three different scenarios.

## 4. Conclusion

Our study describes the interrelationship between the yield of rice and loss through pest species and effective ways to increase the annual yield by controlling insects and improving growth conditions. First, a mathematical model of a rice-pest dynamic system and how the pests harm the annual production of rice is set out, and the model is verified by extensive analysis including the stability at each of the equilibrium points. The annual production of rice decreases with an increasing number of rice pests and vice versa. The rice-pest system is examined through studying transcritical bifurcations, which indicate stability when the number of pests is within acceptable thresholds and instability for above or below the acceptable thresholds.

An optimal control (minimization) problem is newly setup, considering cultural and chemical controls as two control variables to minimize the density of rice pests and the annual losses of rice simultaneously. We conducted analysis to achieve model vigour and obtained necessary conditions for optimality of the problem to determine optimal control variables. The change in the annual production and paddy-loss due to implementing either one or both controls has been compared in detail with the annual production and paddy-loss without control.

Chemical controls, however, are potentially toxic and unsafe with detrimental effects on nearby ecosystems. Therefore, it is recommended to apply chemical control only in emergency situations. We have developed a decision model to determine when pesticides should be applied. The decision model is activated only when the level of insects exceeds the acceptable threshold and the model becomes inactive only when the level of acceptable damage threshold reaches the maximum level, which means that the model becomes inactive when environmental pollution reaches the most acceptable level. This situation continues to recur until the end of cultivation. The acceptable thresholds of pest population can vary with the pest species. In this case, the pesticide application time can change for the change in the acceptable thresholds.

Cultural methods have the potential to control rice pests in agriculture sustainably and simultaneously increase the annual production of rice as shown in Fig. 6. However, the use of chemical controls can further improve the yearly rice yield as shown in Fig. 7. Comparing Fig. 6 and Fig. 7, chemical controls show to be more effective and easier to apply than using cultural controls, contributing to higher annual yield thus contributing to local and global food security. However, when cultural methods are used widely and until the end of cultivation with chemical controls being also applied in emergencies, the system can reduce the growth rate of rice pests by about 33.02% which reduces the annual losses by about one-third, as represented in Fig. 8. We also carried a comparison among the presented three scenarios displayed in Fig. 9. The result of the comparison finds that if cultural methods can be used till the end of cultivation and chemical controls can be applied only in emergencies, it would be the best control strategy for increasing the annual rice production and controlling rice pests.

This study could be helpful in agriculture by enriching rice production and controlling pests naturally and artificially. However, the long-term effects of using chemicals even in smaller numbers is an outstanding issue – chemical control can bring short-term relief but may damage the ecosystem with time. Here, the inclusion of changing water quality of nearby aquatic ecosystems, such as rivers or canals, in the mathematical model as a feedback mechanism could be considered. Furthermore, the introduction of biological controls such as certain predators or parasites as alternatives to chemical controls would make sense to be studied in combination to have more control variables and a decision model aimed at increasing simultaneously productivity and sustainability.

## Author contribution

Author contributions have been evaluated according to CRediT.

**Sajib Mandal**: Conceptualisation, Methodology, Software, Validation, Formal Analysis, Investigation, Data Curation, Writing - Original Draft, Writing – Review & Editing, Visualisation.

**Sebastian Oberst:** Conceptualisation, Methodology, Investigation, Validation, Writing - Review & Editing. **Md. Haider Ali Biswas:** Conceptualisation, Methodology, Validation, Resources, Supervision, Project administration.

**Md. Sirajul Islam:** Conceptualisation, Supervision.

## Supplementary Materials

### Supplementary A. Modelling of the rice-pest dynamic system and its biological control

#### Supplementary Material A.1 Study area

This study has been conducted globally without being confined to a specific region because food security and pest management are global issues. It is not possible to reduce the annual loss of rice and control pest infestation by adopting certain control strategies in certain areas of a country. Here, we consider two control strategies, cultural methods, and chemical control, to increase the annual production of rice by controlling the pests of rice in the field. Cultural methods which consist of natural controls such as soil rotation, crop variation, and natural enemies are used as the first and foremost control strategy because of having natural capacity of increasing the production of rice and controlling rice pests. On the other hand, chemical controls consisting of various pesticides such as neonicotinoids, glyphosate, DDT, BHC, and 2,4-D are used only for emergencies because of having adverse effects on the environment and crop quality. Here, we do not define any specific pesticide because pesticides can vary from pest to pest.

#### Supplementary Material A.2

Lotka-Volterra Model in an Optimal Control Problem Setting

The general form of the Lotka-Volterra model can be written in terms of a pair of autonomous differential equations

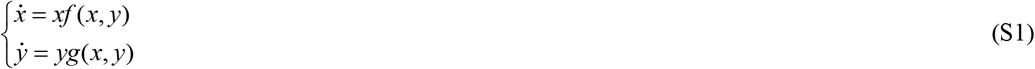

with *x* and *y* being the prey and predator populations and *f*(*x, y*) and *g*(*x, y*) being the species’ growth rates.

An optimal control problem (OCP) is concerned with the state and the control variable. Consider *x*(*t*) and *u*(*t*) are these two variables of an OCP respectively, then *x*(*t*) satisfies

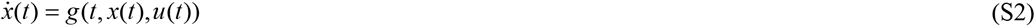

with *g* being continuously differentiable. The objective of this OCP is to find a piecewise continuous control function *u*(*t*) and its corresponding state variable *x*(*t*) to optimize (either maximize or minimize) a specific objective function [31]. An OCP can be described in the following way

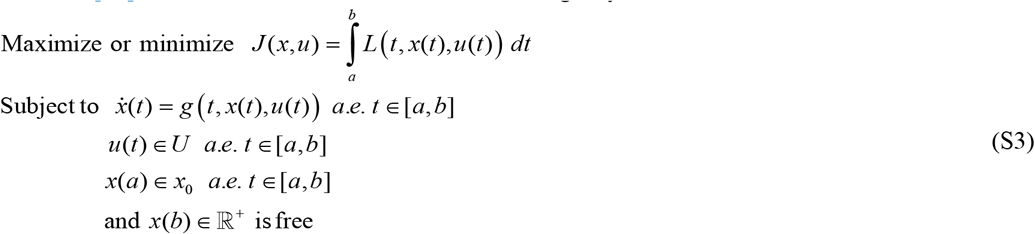

with *u*(*t*) and *x*(*t*) both being piecewise *C*^1^ differentiable with *t* in [*a, b*] being the time interval where 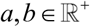 and *a < b*. In the problem, *u*(*t*) belongs to a certain space *U* that may be a piecewise continuous function or a space of measurable function which satisfies all the constraints of the problem. Therefore, (*x*\*, *u*\*) constitutes the optimal solution of the OCP in case the costs can be minimized overall admissible processes [31].

#### Supplementary Material A.3 Assumption and model formulation of the rice-pest dynamic system

Let *x*_1_(*t*) be the annual production of rice and *x*_2_(*t*) be the corresponding pest population at a time *t* where the annual production of rice acts as the prey population, and the corresponding pest population acts as the predator population. In this regard, the pest population consumes and damages the rice growth which declines the annual production of rice. The relationships between the production of rice and the density of pests are as follows [24,34]:

i. At the initial state, let the reproduction rate of rice be *α*_1_. For the consumption of rice by the pest population, a part of the annual production of rice is lost which let’s *β*_1_ be the loss rate of *x*_1_(*t*) due to the consumption of *x*_2_(*t*). Then the reproduction rate of rice can be represented by

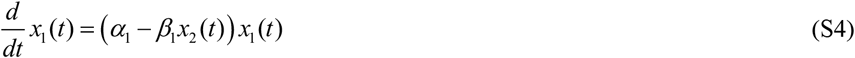

where *β*_1_*x*_1_(*t*) *x*_2_(*t*) presents the annual loss of rice due to pests.
ii. On the other hand, the pest population in the system depends on rice for feeding. Therefore, the pest population proportionally increases with the increasing rice production. Let *β*_2_ be the energy gain rate of pest population by consuming rice. Again, since the pest population depends on the production of rice for feeding, their growth and population number will be reduced when the production of rice declines. Therefore, *α*_2_ is taken as the decline rate of the pest population proportionally with the decline of rice production. Then the growth rate of the pest population can be represented by

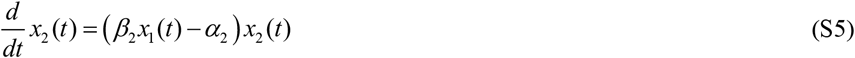

where *β*_2_*x*_1_(*t*) *x*_2_(*t*) presents the annual consumption of rice by the pest population.

The rice-pest system can be written in terms of a pair of Nonlinear Ordinary Differential Equations (NODEs) in form of a Lotka-Volterra model:

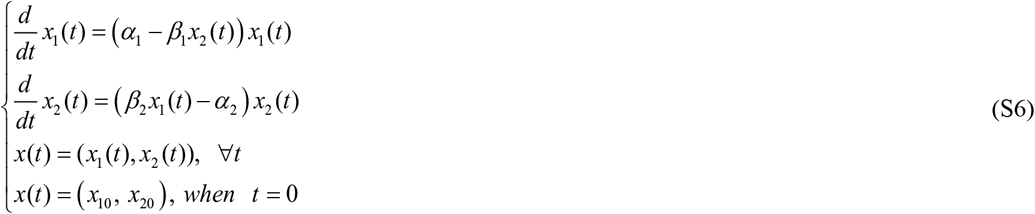

Where *x*_1_(*t*) and *x*_2_(*t*) respectively present the annual production of rice and the density of the rice pests at time *t*, and *x*_10_ and, *x*_20_ respectively show the initial conditions of *x*_1_(*t*) and *x*_2_(*t*) i.e. the values of *x_2_(t*) and *x*_2_ (*t*) at *t* = 0.

The production rate of rice does not damage only for the attack of pests, but also it may be damaged owing to environmental impacts such as drought, global warming, etc. Similarly, the pest population may be reduced due to natural causes such as floods, droughts, etc. Therefore, two parameters i.e., *d*_1_ (decrease rate of rice production due to natural causes not related to the pest population), and *d*_2_ (death rate of pest population due to natural causes not related to the deficiency of rice) need to be considered in the model as well:

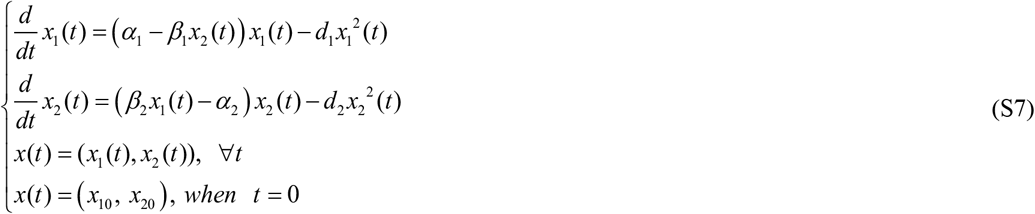

Here, the term 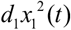 presents the decrease rate due to intraspecific competition in species *x*_1_(*t*) due to natural causes that are not related to *x*_2_(*t*) e.g., viral infections, droughts, or floods [35,36]. Similarly, the term 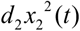 shows the intraspecific competition between *x*_2_ (*t*) due to natural causes that are not related to *x*_2_(*t*), e.g., viral infection and heavy rains [35,37].

A schematic diagram is shown in Fig. S1 which represents the dynamic relations of this model.

**Fig. S1.**
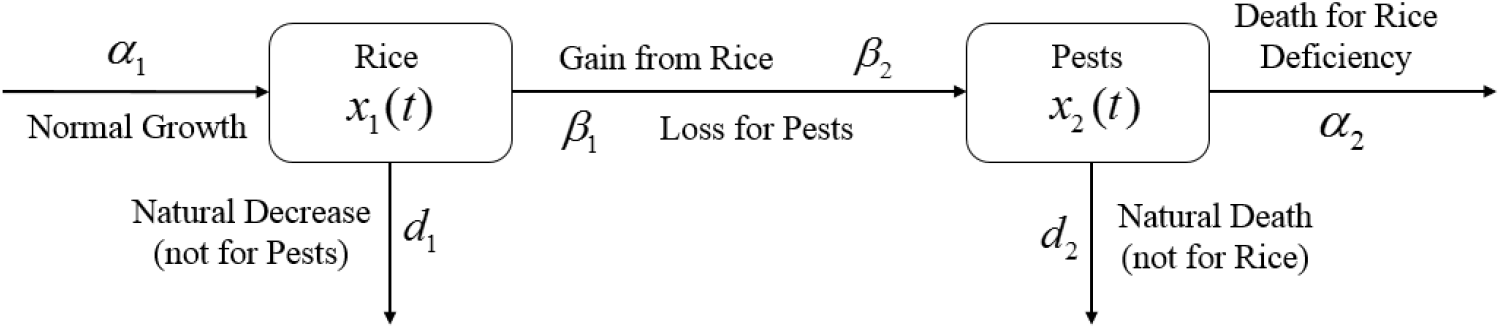
A schematic diagram of the rice-pest system (S7) disclosing the interrelationship between rice and corresponding rice pests along with the impact of adverse environment.

#### Supplementary Material A.4 Positivity Analysis

In the following, we intend to show that the rice-pest system (S7) is bounded in a positive region and the dynamic species of the system (S7) consist of non-negative values at any time [38].

##### Theorem A.1.

*The rice-pest system (S7) is bounded in a positive region* 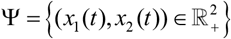.

**Proof:** Consider *N*(*t*) = *x*_1_(*t*) + *x*_2_(*t*) and 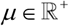, then we can write

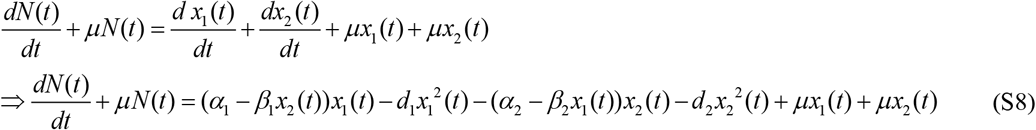

Since the consumption of pests and the losses of rice are approximately equal at the equilibrium point i.e., *β*_1_ ≈ *β*_2_, Eq. (S8) takes the following form

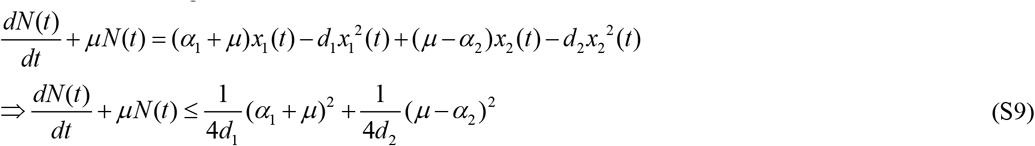

Applying the principle of differential inequalities in Eq. (S9), we get

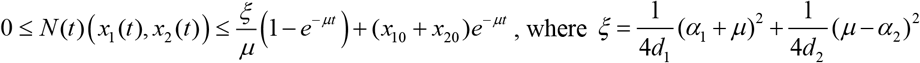

Taking the limit as *t* → ∞ we obtain 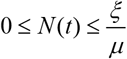, hence the region of attraction for the system can be formulated as 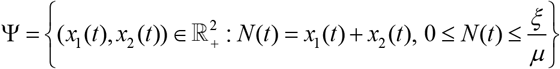 with *ξ*, 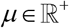. This indicates that the system is positively bounded in Ψ.

##### Theorem A.2.

*Each species of the model (S7) contains a non-negative real value for all t* ≠ 0.

**Proof:** To estimate the solution, consider the first equation of the model (S7) given as

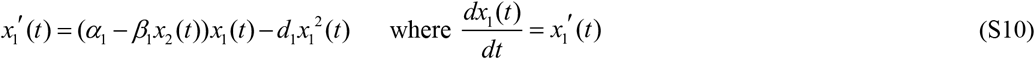

According to the condition of positivity, Eq. (S10) takes the following form

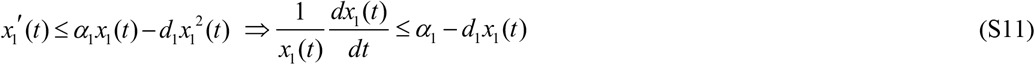

After solving Eq. (S11), the solution of *x*_1_(*t*) becomes 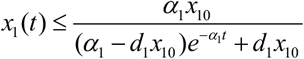. Therefore, 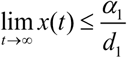 i.e. 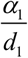 is the upper limit of *x*_1_(*t*) which implies that 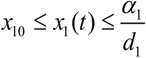 i.e., *x*_1_(*t*) is non-negatively bounded. Similarly, it can be shown that 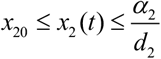. Hence the species of the system are positively bounded.

#### Supplementary Material A.5 Equilibrium Points

To conduct the stability analysis of the rice-pest system (S7), we first obtain the equilibria of the system, considering three different situations [38]:

i. The *trivial* or *pre-cultivation* case refers to the situation prior to cultivation. No plants and pests exist yet. Therefore, the pre-cultivating equilibrium point is (*x*_10_, *x*_20_) = (0, 0).
ii. The *pest-free* case refers to the pre-existing state of rice pests, indicating that there are rice plants, but no pests. Here, the pest-free equilibrium point is 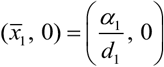.
iii. A *co-existence* or *farming* situation refers to a competitive case where both rice and pests coexist. The co-existing equilibrium point of the rice-pest system (S7) is

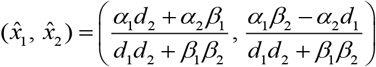

#### Supplementary Material A.6

Stability Analysis

To illustrate the nature of the equilibrium points of the model (S7), a stability analysis has been conducted. In this case, let’s consider system (S7) in the following vectorial form [38]

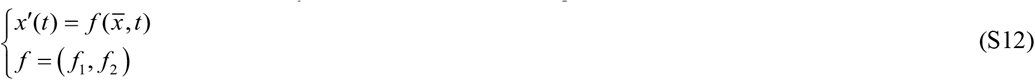

where 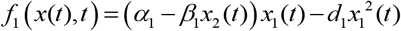 and 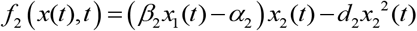

After evaluating the vector field, the Jacobian matrix is obtained as the following

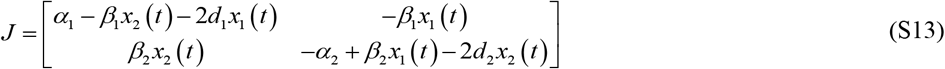

##### Theorem A.3.

*System (S7) approaches a saddle-node at the pre-cultivation equilibrium*.

**Proof:** To prove the theorem, evaluate the Jacobian matrix (S13) at (*x*_10_, *x*_20_) which becomes of the following form

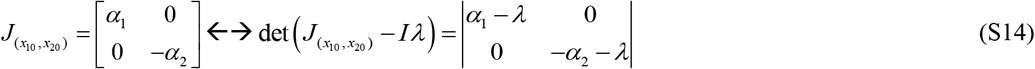

The characteristic equation of Eq. (S14) is

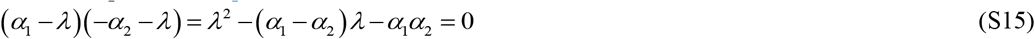

Hence the eigenvalues of Eq. (S15) are *λ*_1_ = *α*_1_ and *λ*_2_ = –*α*_2_. Since the eigenvalues are real and are opposite in sign, the *pre-cultivating equilibrium* represents a saddle point.

For an autonomous system, an equilibrium point is a saddle point if the characteristic equation consists of opposite signed (one positive and one negative) real eigenvalues [39]. A stable point exhibits a mean position between the stability and instability of a point. For a biological example in two dimensions, the dynamic species appear to have reached a stable equilibrium situation but, their trajectory bends to the other side instead.

##### Theorem A.4.

*The system described by Eq. (S7) at pest-free equilibrium approaches a saddle point*.

**Proof:** To test the stability at *pest-free equilibrium*, evaluate the Jacobian matrix (S13) at 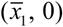 through its characteristic equation by calculating det 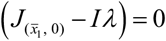

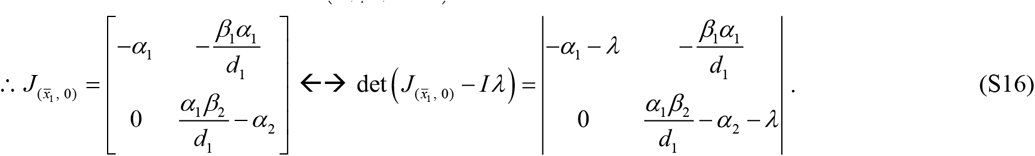

Therefore, the characteristic equation of Eq. (S16) is

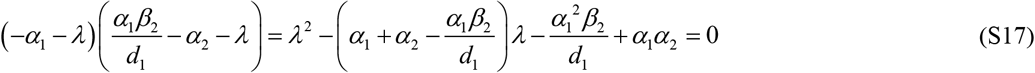

Similarly, the eigenvalues of Eq. (S17) become *λ*_1_ = – *α*_1_, and 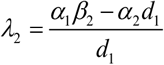 with *λ*_2_ being strictly positive since *α*_1_*β*_2_ > *α*_1_*d*_1_ which indicates that the *pest-free equilibrium* represents a saddle point.

##### Theorem A.5.

*The rice-pest system (S7) approaches a spiral node at the co-existing equilibrium point*.

**Proof:** Evaluating the Jacobian matrix (S13) at the co-existing equilibrium point, it takes the following form

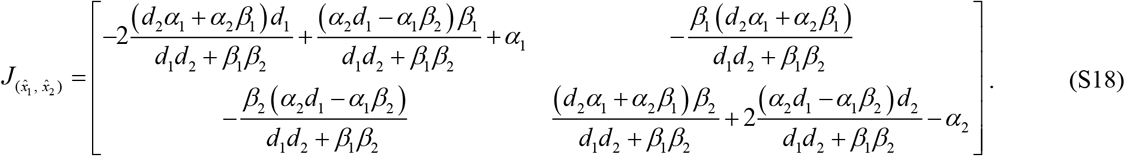

The characteristic equation of Eq. (S18) is again calculated via 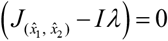

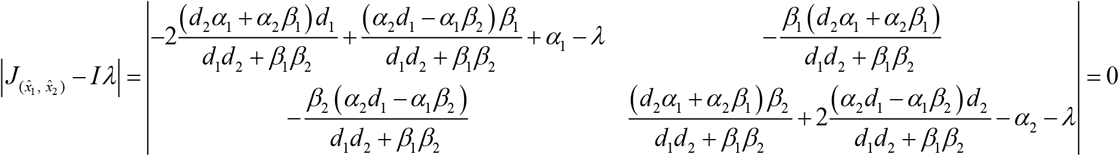

with the eigenvalues becoming

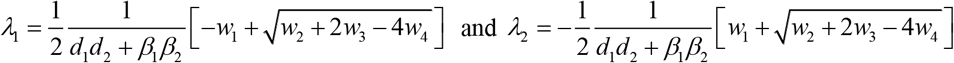

with *w*_1_ = *α*_1_*d*_2_ (*β*_2_ + *d*_1_) + *d*_1_ *α*_2_ (*β*_1_ – *d*_2_), 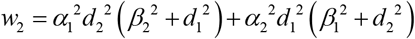, *w*_3_ = *d*_1_*d*_2_(*α*_1_*α*_2_*d*_1_(*β*_1_ + *d*_2_) + *α*_2_*β*_1_(*α*_2_*d*_1_ + *α*_1_*b*_2_)–*α*_1_*b*_2_*d*_2_(*α*_1_ + *α*_2_), and *w*_4_ = *β*_1_*β*_2_(*α*_1_*α*_2_*β*_1_*β*_2_ + *d*_2_*α*_1_^2^*β*_2_–*d*_1_*α*_2_^2^*β*_1_)

Both the eigenvalues are complex numbers which indicate that the *co-existing equilibrium* approaches a spiral node.

**Fig. S2.**
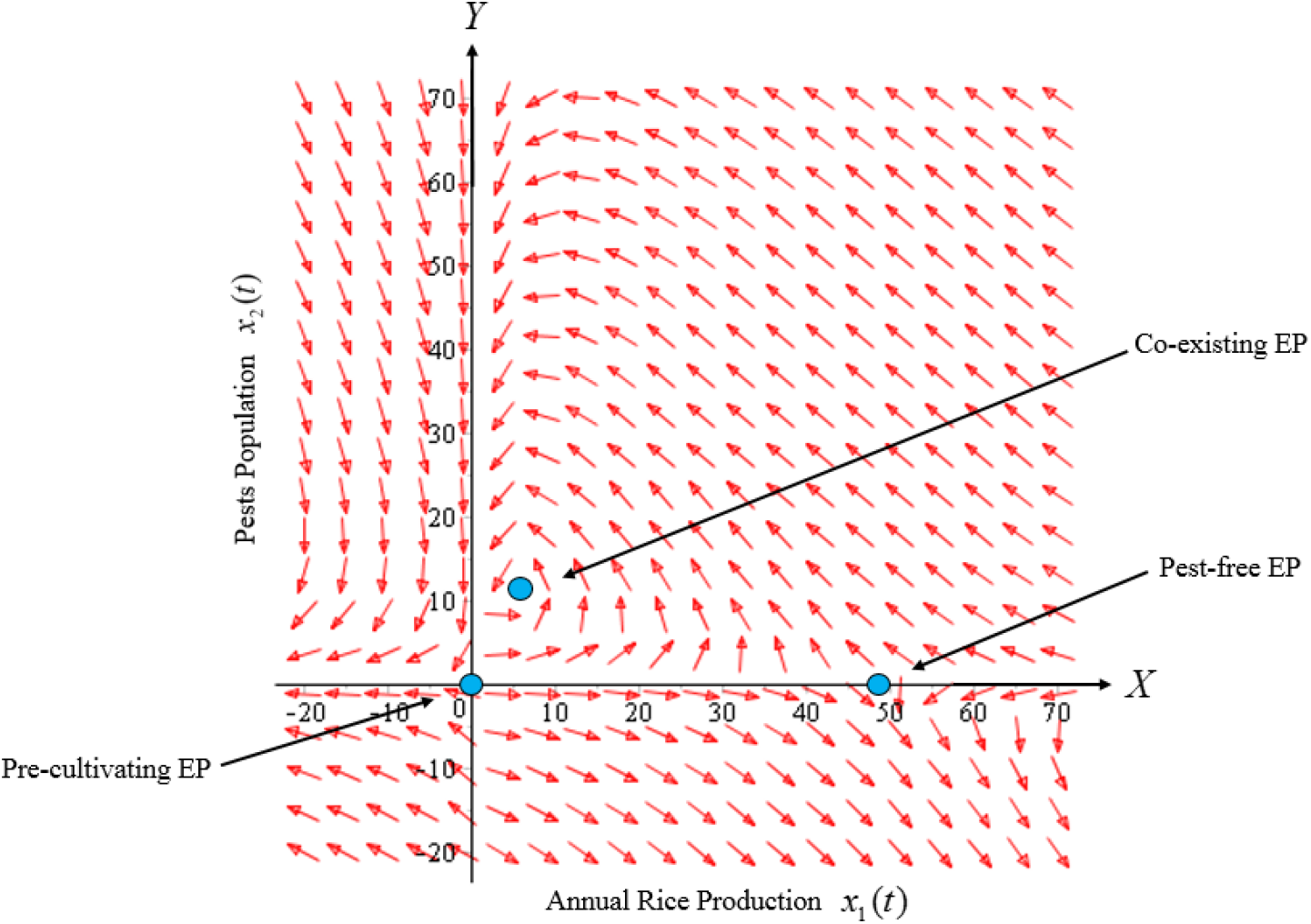
Phase portrait discloses the characteristics of three equilibrium points (EP) of the rice-pest system (S7). Here, the co-existing equilibrium point is a spiral node, and the others, pre-cultivating and pest-free equilibrium points, are saddle points.

### Supplementary Material B. Bifurcation analysis for the rice-pest system

In this section, we investigate the rice-pest system (S7) through transcritical bifurcation analysis [40]. For a transcritical bifurcation exists a non-destructible fixed point over the whole bifurcation parameter range, which, however, changes its stability characteristic for altered bifurcation parameter values [40]. For transcritical bifurcation analysis of rice-pest system (S7), it is more convenient to make the rice-pest system (S7) dimensionless to reduce the number of parameters [40]. In this case, we introduce the dimensionless variables 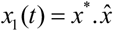 and 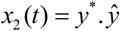 and consider the dimensionless time *t* = *τ*^*^*σ*. Removing the symbols ‘*’ and ‘^’, the system (S7) becomes dimensionless given as

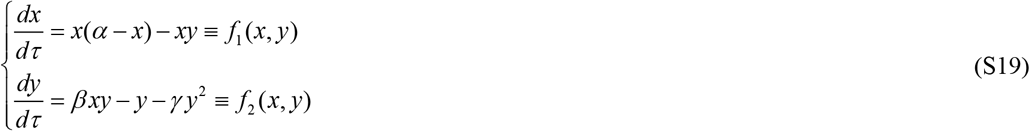

where 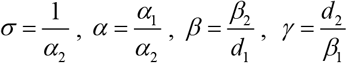.

The biologically meaningful equilibria of the system (S19) are the non-negative solutions of *f*_1_(*x, y*) = 0 and *f*_2_(*x, y*) = 0. The rice (prey) isocline consists of the axis *x* = 0 and the straight line *y* = *α*–*x* and the pests (predator) isocline consists of the axis *y* = 0 and the line 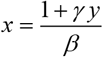 [40]. Correspondingly, the system (S19) is bounded by two equilibrium points, the trivial/pre-cultural equilibrium *E*_0_ (*x, y*) = (0,0) and pest-free equilibrium *E*_1_(*x, y*) = (*α*, 0). Besides, an interior equilibrium *E*_*_ (*x, y*) = (*x_*_, y_*_*) where 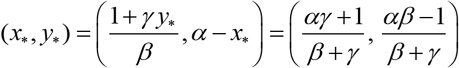 can be found at the intersection of the two isoclines. For the existence of *E*_*_, the value of the parameters must follow the conditions *αγ* + 1 ≥ 0, *β* + *γ* > 0 and *αβ* – 1 ≥ 0.

#### Theorem B.1.

*The system (S19) experiences transcritical bifurcation at the pest-free equilibrium point E*_1_ (*α*, 0) *as the growth parameter α passes through the critical value α*^*^.

**Proof:** At the pest-free equilibrium point *E*_1_(*α*, 0), the associated Jacobian matrix of the system (S19) takes the form:

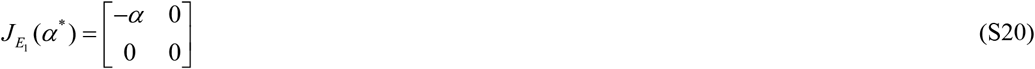

The set of eigenvalues of *J*_*E*_1__ (*α*^*^) is 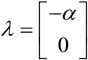 i.e. one eigenvalue is zero and the other is negative since *α* > 0.

Therefore, to examine the nature of the system at *E*_1_, we have applied *Sotomayor’s theorem* [41]. For this purpose, we consider the system (S19) as

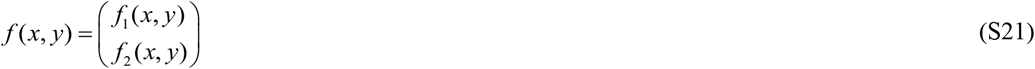

Let the eigenvectors corresponding to the zero eigenvalues of *J*_*E*_1__ (*α*^*^) and *J*_*E*_1__^*T*^(*α*^*^) be *V* and *W*, respectively, where 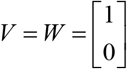. From Eq. (S21), we have 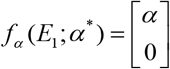, 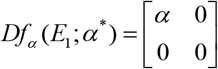 and 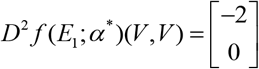. Here, *Df* denotes the partial derivative of *f* with respect to *x* and *y*, and *Df_a_* denotes the partial derivative of *f* with respect to the parameter *α*. Therefore,

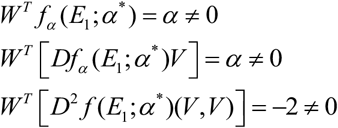

Hence, there is a saddle-node bifurcation at the nonhyperbolic equilibrium point *E*_1_(*α*, 0) at the bifurcation value *α*. For *α* < 0, there is no equilibrium point. For *α* = 0, the *f*_1_(*x, y*) = –*x*^2^ is structurally unstable and the bifurcation value *α* = 0. Therefore, there is a transcritical bifurcation at the origin for *α* = 0. There are two equilibria at origin (0,0) and *E*_1_ (*α*,0) [41].

#### Theorem B.2.

*The system (S19) experiences transcritical bifurcation at the equilibrium point E*_*_ (*x_*_, y_*_*) *as the growth parameter of pest population β passes through the critical value β*^*^.

**Proof:** At the equilibrium point *E*_*_(*x_*_, y*_*_), the associated Jacobian matrix of the system (S19) takes the form:

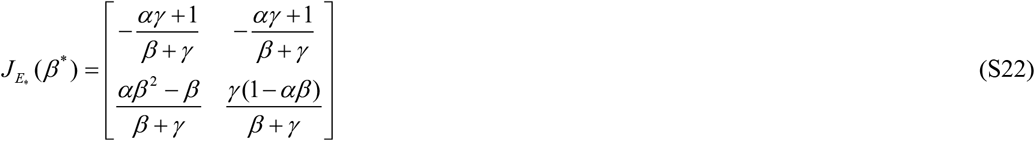

It can be shown that one eigenvalue of *J*_*E*\*_ (*β*^*^) is negative and the other is zero for the condition *αβ* = 1 [41]. To investigate the nature of the system at *E*_*_, we have applied *Sotomayor’s theorem* [41]. Let the eigenvectors correspond to the zero eigenvalues of *J_E*_*(*β*^*^) and *J_E*_^T^*(*β*^*^) be *V* and *W*, respectively, where 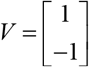 and 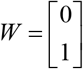. From Eq. (S21), we have 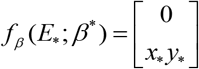, 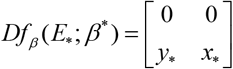 and 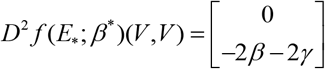. Therefore,

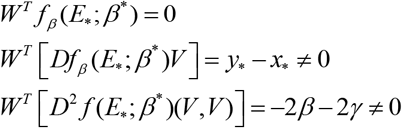

Hence the system (S19) satisfies all the necessary conditions of *Sotomayor’s theorem* and thus the system (S19) experiences a transcritical bifurcation at the coexistence equilibrium point *E*_*_, for the bifurcation parameter *β* [41].

#### Supplementary Material B.1 Significance of β as a transcritical bifurcation paremeter

We have numerically investigated the dynamic behaviour of the system (S19) for the variation in the growth of pest populations (*β*). Let *β*_0_ be the initial condition for the existence of *E*_*_. The parameters must follow the conditions *αγ* +1≥0, *β* + *γ* > 0 and *αβ* – 1 ≥ 0 and to estimate *β*_0_, we consider *α* = 1 and *γ* = 0.001 which satisfy all the conditions. Therefore, we get *β*_0_ = 1 after calculating det(*J_E*_* (*β*^*^)) = 0 [42].

For *β* < 1, there is no intersection between the rice isocline and pests isocline as represented in Fig. S3(a). In this case, the system (S19) experiences unstable and no equilibrium point. For *β* = 1, the isoclines of rice and pests intersect at the pest-free equilibrium point *E* (1,0) as shown in Fig. S3(b). For *β* > 1, the system (S19) experiences an interior point *E_*_*(*x_*_*,*y_*_*) between the origin (0, 0) and *β*(1,0) as represented in Fig. S3(c). The nature of the system (S19) is illustrated at the interior equilibrium point *E_*_* (*x_*_, y_*_*) for the different values of *β* > 1 which is described in Fig. S4. The rice-pest system (S19) is stable at the interior critical point *E* for the bifurcation parameter *β* ∈ [1,30]; e.g., the system is stable for choices of *β* = 2, *β* = 8, and *β* = 30 as shown in Fig. S4(a)-4S(c), respectively. But for *β* = 31, there is a limit cycle at *E_*_* which describes that the system (S19) remains in oscillation with respect to time and becomes unstable as represented Fig. S4(d). Hence, the maximum value of *β* is *β^*^* = 30. Moreover, the transcritical bifurcation diagram with respect to the bifurcation parameter *β* is represented in Fig. S5. The diagram also examines that the rice-pest system (S19) is stable only for *β* ∈ [1,30] and gradually tends to an imbalance situation with the increase of the growth of the pest populations (*β*) and even the system will be defeated for high growth rate (*β* > 30).

Hence the bifurcation analysis suggests to control the pest population for the sustainable management of the rice-pest system (S7), otherwise, the high pest populations will destroy the system.

**Fig. S3.**
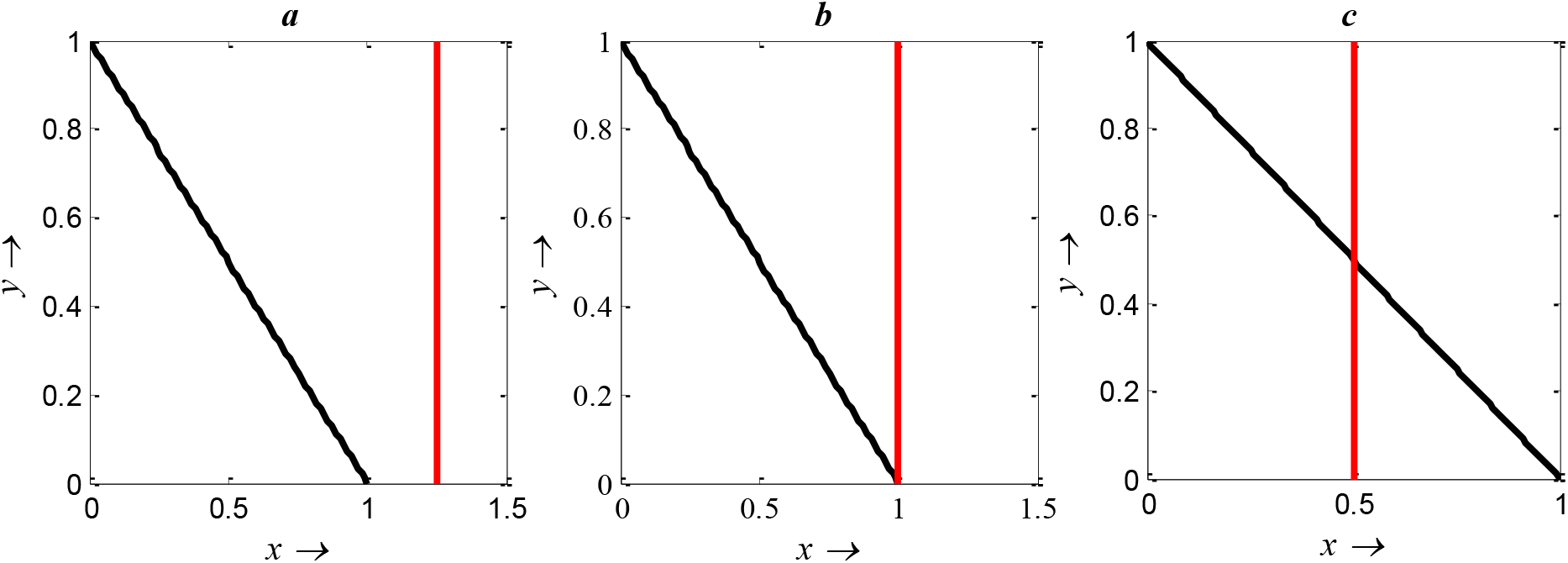
Diagram for the isocline of the rice (vertical) and the pest’s isocline (inclined): (a) *α* = 1, *β* = 0.8, *γ* = 0.001; (b) *α* = 1, *β* = 1, *γ* = 0.001; (c) *α* = 1, *β* = 2, *γ* = 0.001.

**Fig. S4.**
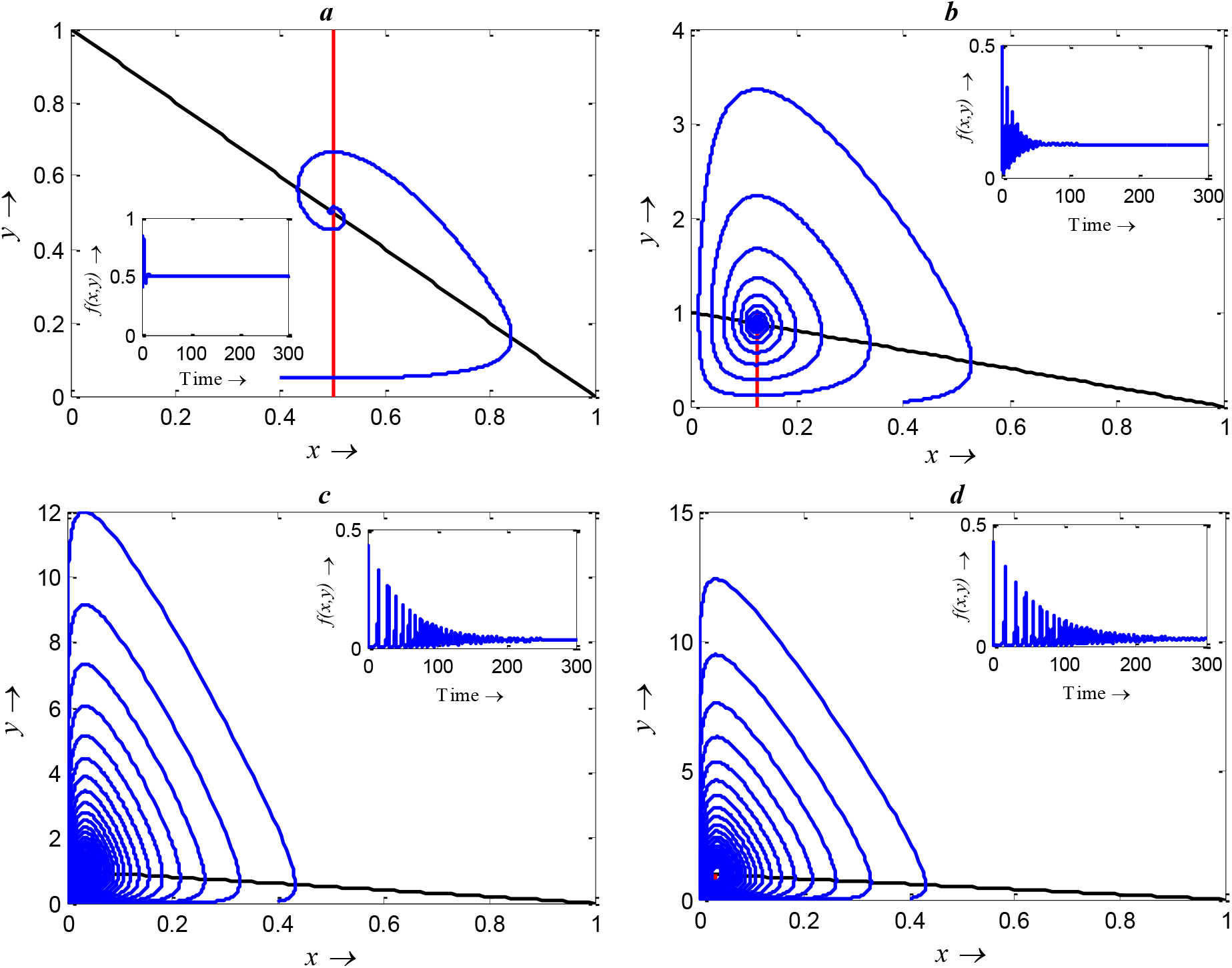
Phase plane describing the nature of the system (S19) at the interior point for the variation of *β*: (a) *β* = 2, (b) *β* = 8, (c) *β* = 30 and (d) *β* = 31, where *α* = 1 and *γ* = 0.001 remain same.

**Fig. S5.**
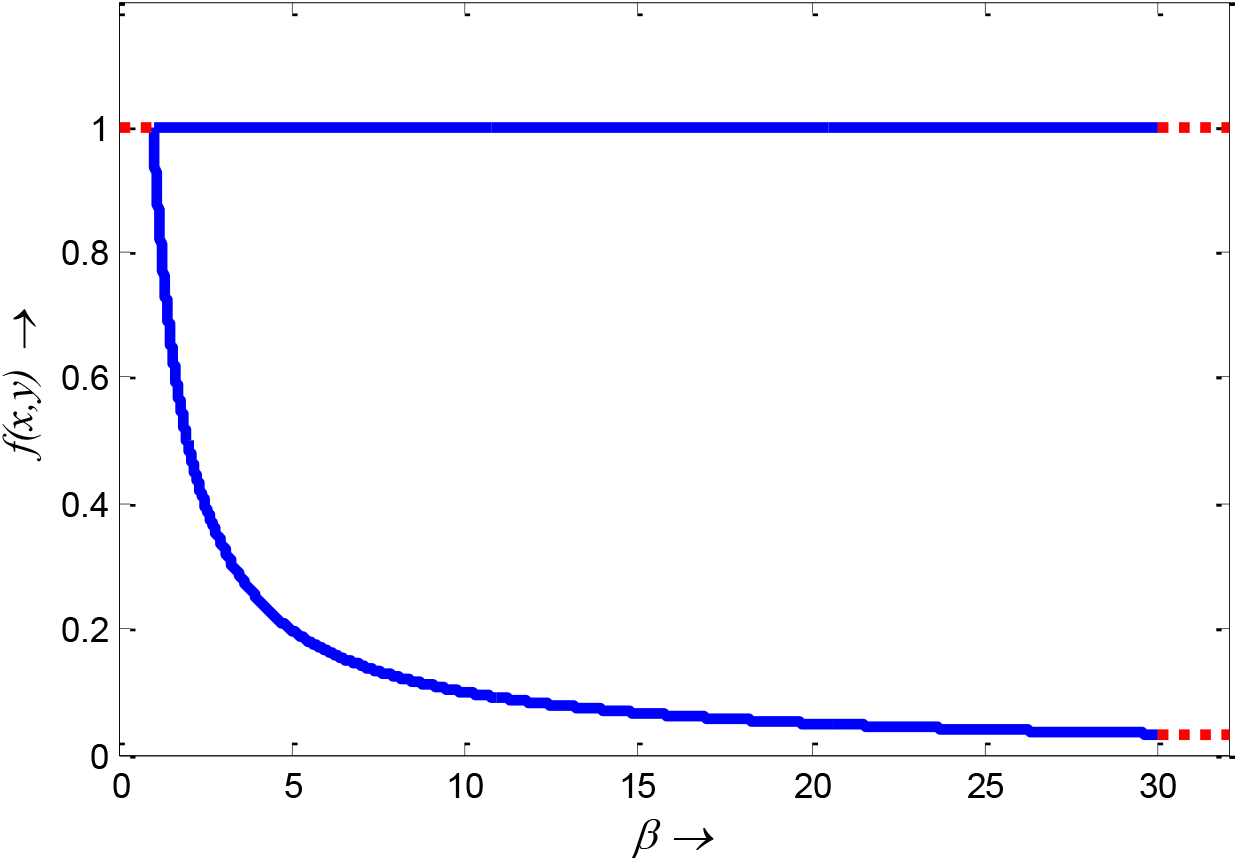
Transcritical bifurcation diagram of the rice-pest system (S19) for the bifurcation parameter *β* showing the system (S19) stable for *β* ∈ [1,30] (continuous blue line) and unstable for 0 ≤ *β* < 1 and *β* > 30 (dash red line). The bifurcation diagram reveals that the rice-pest system (S19) is present within the acceptable thresholds of the pests population and is destroyed for above and below the acceptable thresholds.

### Supplementary Material C. Data Analysis

All the parameters are secondary (derived/estimated or collected from other sources), two of which are estimated. For parametric estimation, we first studied deeply the rice-pest system. Then we conducted statistical analysis for parametric estimation after collecting and observing the corresponding data collected from different sources [26,32,33]. The estimation of parameters used in this study is described below.

#### *Estimation of α*_1_

**Table A1.** Annual data of the production of rice [26]

| Data\Years | 2018 | 2013 | 2008 | 2003 | 1998 | 1993 | 1988 | 1983 | 1978 | 1973 | 1968 | 1963 |
| --- | --- | --- | --- | --- | --- | --- | --- | --- | --- | --- | --- | --- |
| Harvested area (MH) | 167.12 | 165.12 | 160.08 | 148.45 | 151.68 | 146.45 | 146.40 | 142.83 | 143.50 | 136.57 | 129.26 | 120.15 |
| Production (MMT) | 782.00 | 742.50 | 687.05 | 586.93 | 578.81 | 529.60 | 487.46 | 448.02 | 385.21 | 334.93 | 288.62 | 247.12 |
| Production rate (metric tons/ha) | 4.679*<br>($\alpha_1$) | 4.497 | 4.292 | 3.954 | 3.816 | 3.616 | 3.330 | 3.137 | 2.684 | 2.452 | 2.286 | 2.057 |

Here, MH presents Millions of hectares, and MMT means Millions of metric tons.

Therefore, the production rate of rice is *α*_1_ = 4.679 metric tons/ hectare

#### *Estimation of α*_2_

**Table A2.**
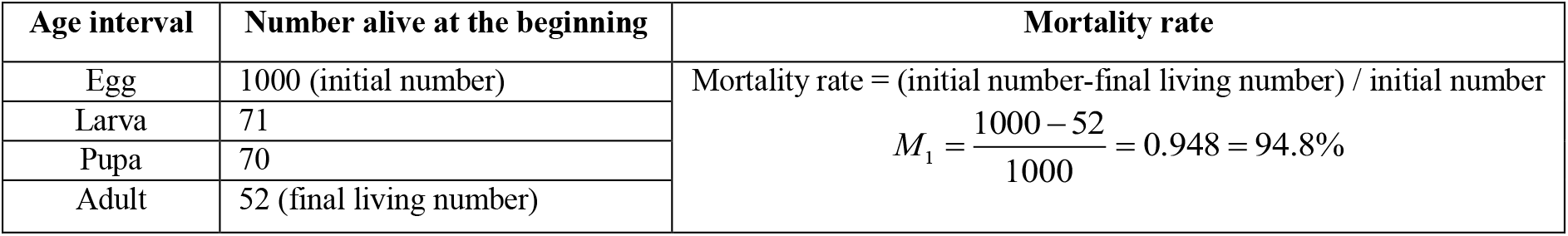
Life table for *Heliothis zea* in sweet corn 10 October 1976 – 20 November 1976 [33]

**Table A3.**
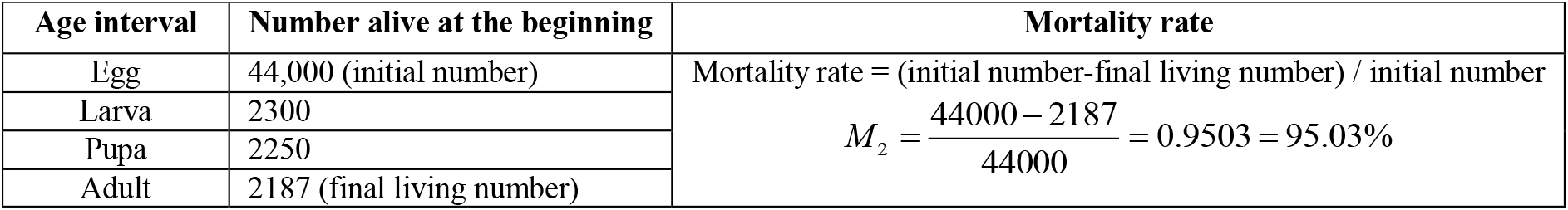
Cohort life table from a hypothetical caddisfly population [32].

The average mortality rate is 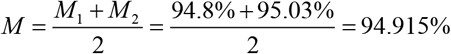

∴*α*_2_ = 94.915%

#### *Estimation of β*_1_ *and β*_2_

Annual losing rate of rice by pests is about 37% =37/100= 0.37 [6,7,9]. Therefore *β*_1_ = 37% or 0.37. Since the predation rate of predators is approximately equal to the loss rate of prey population in co-existing equilibrium [6,7,9], therefore *β*_2_ = *β*_1_ = 37% or 0.37.

#### *Estimation of d*_1_ *and d*_2_

The natural death rate of paddy plants (without pests) is approximately 10% (on average) [26]. Therefore *d*_1_ = 10% or 0.10

The death rate of pests due to natural causes or chemical control = (1 – normal mortality rate of pests) where 1 is the total probability index.

Therefore, *d*_2_ = (1 - *α*_2_) = (1 - 0.94915) = 0.05085 = 5.085%

The parameters with numerical values are displayed in Table 1. The parameters are also verified by comparing the results with other research [2,4,8–10,12], employing numerical simulations as deeply described in Section 3.

